# Local haplotype visualization for trait association analysis with crosshap

**DOI:** 10.1101/2023.05.07.539781

**Authors:** Jacob I. Marsh, Jakob Petereit, Brady A. Johnston, Philipp E. Bayer, Cassandria G. Tay Fernandez, Hawlader A. Al-Mamun, Jacqueline Batley, David Edwards

## Abstract

**Summary:** GWAS excels at harnessing dense genomic variant datasets to identify candidate regions responsible for producing a given phenotype. However, GWAS and traditional fine-mapping methods do not provide insight into the complex local landscape of linkage that contains and has been shaped by the causal variant(s). Here, we present ‘crosshap’, an R package that performs robust density-based clustering of variants based on their linkage profiles to capture haplotype structures in a local genomic region of interest. Following this, ‘crosshap’ is equipped with visualization tools for choosing optimal clustering parameters (ε) before producing an intuitive figure that provides an overview of the complex relationships between linked variants, haplotype combinations, phenotypic traits and metadata.

**Availability and implementation:** The ‘crosshap’ package is freely available under the MIT license and can be downloaded directly from CRAN with R>4.0.0. The development version is available on GitHub alongside issue support (https://github.com/jacobimarsh/crosshap). Tutorial vignettes and documentation are available (https://jacobimarsh.github.io/crosshap/).

## 1. Introduction

Rapidly accumulating genomic sequence information presents opportunities to identify and characterize the effects of variants conferring beneficial adaptations. However, with increasing population sizes, the dimensions of variant information increase exponentially, raising interpretability issues when analysing large genomic datasets. Genome-wide association studies (GWAS) offer a scalable approach to identify candidate regions responsible for influencing a phenotype, though they do not provide insight into the complex local landscape of linkage that contains and, in many cases, has been shaped by selection for the causal variant(s).

Statistical fine-mapping is the standard approach for visualizing phenotypic associations for local genomic variants surrounding a GWAS result (Schaid et al., 2018). These methods excel at isolating candidate variants responsible for simple traits with strong phenotypic associations caused by explainable genomic changes such as nonsense mutations (Pruim et al., 2010). However, fine-mapping only facilitates users to evaluate variants in a region of interest by unidimensional phenotypic association scores, with limited tools to explore linkage relationships between variants (Marsh et al., 2022). In addition, fine-mapping tools do not link analysis of genomic variation with biological or geographical features of individuals, such as level of domestication or region collected, which may be important co-factors influencing phenotypic association scores.

Local haplotyping defines groups of individuals based on allelic combinations across loci in a region of interest (Belzile et al., 2020). The primary advantage of this approach over statistical fine-mapping is that it connects shared genomic variation to the individual, which aids in capturing the phenotypic effects of haplotypes, and allows for characterization of the identified populations that possess shared allelic combinations within the region of interest (Marsh et al., 2022). The approach used by RFGB v2.0 (Wang et al., 2019) and CandiHap (Li et al., 2023) for capturing local haplotypes considers unique combinations of all SNPs in a window within or centred on a gene of interest. However, across a wider interval with dozens, hundreds or thousands of variant loci, minor divergences between individuals will result in a rapid expansion of the number of unique haplotypes identified, which restricts effective analysis between groups.

LD-based local haplotyping approaches such as HaplotypeMiner (Tardivel et al., 2019) enable analysis of wider windows through variant dimension reduction, with ‘tagging’ to prune redundant SNPs in high LD before defining haplotypes. Alternatively, HapFM (Wu et al., 2022) is an end-to-end causal candidate haplotype identification tool that clusters individuals into haplotypes from local variation surrounding a GWAS result. The strengths of HaplotypeMiner and HapFM lie in their ability to synthesize general patterns of linkage from large complex variant data to assign individuals into relevant groups for downstream analysis. However, current local haplotyping software is limited in its ability to convey the relevance of haplotyping results to the user. Realising the full potential of LD-based local haplotyping requires drawing connections between the groups of linked variants/haplotype combinations identified and meaningful phenotype and metadata differences between individuals. Informative genomic visualizations are needed to de-convolute complex linkage structures by displaying the relationships between variants within and between each marker group whilst also connecting both marker group and haplotype combination results back to individual-level differences.

Here we present ‘crosshap’, which performs robust density-based clustering of variants based on their linkage profiles to capture haplotype structures in local genomic regions. ‘crosshap’ significantly improves on current software by supporting highly transparent haplotyping, equipping the user with a wide range of tools to understand patterns of local genomic variation through curated visualizations displaying raw data points. Visualization tools are provided by ‘crosshap’ for choosing optimal clustering parameters (ε) and producing intuitive ‘crosshap’ figures that present information on the complex relationships between linked variants, haplotype combinations, and phenotypic/metadata traits of individuals.

## 2. Materials and Methods

### 2.1 Inputs

The ‘crosshap’ workflow is outlined in Figure 1. For haplotyping, ‘crosshap’ requires genomic variant information in VCF format, delimited to a region of interest (see https://jacobimarsh.github.io/crosshap/articles/Delimiting_region.html), along with a square matrix reporting a pairwise LD statistic (e.g. R^2^) between all loci. Phenotype and metadata are two additional input classes describing features of the individuals in the VCF that aid visualization and interpretation of haplotyping results. Phenotype supports any numerical description of individuals, such as yield, height or time to maturity. Metadata, which is optional input, supports any categorical description of individuals, such as level of domestication or the geographic region accessions were sampled from.

**Figure 1.**
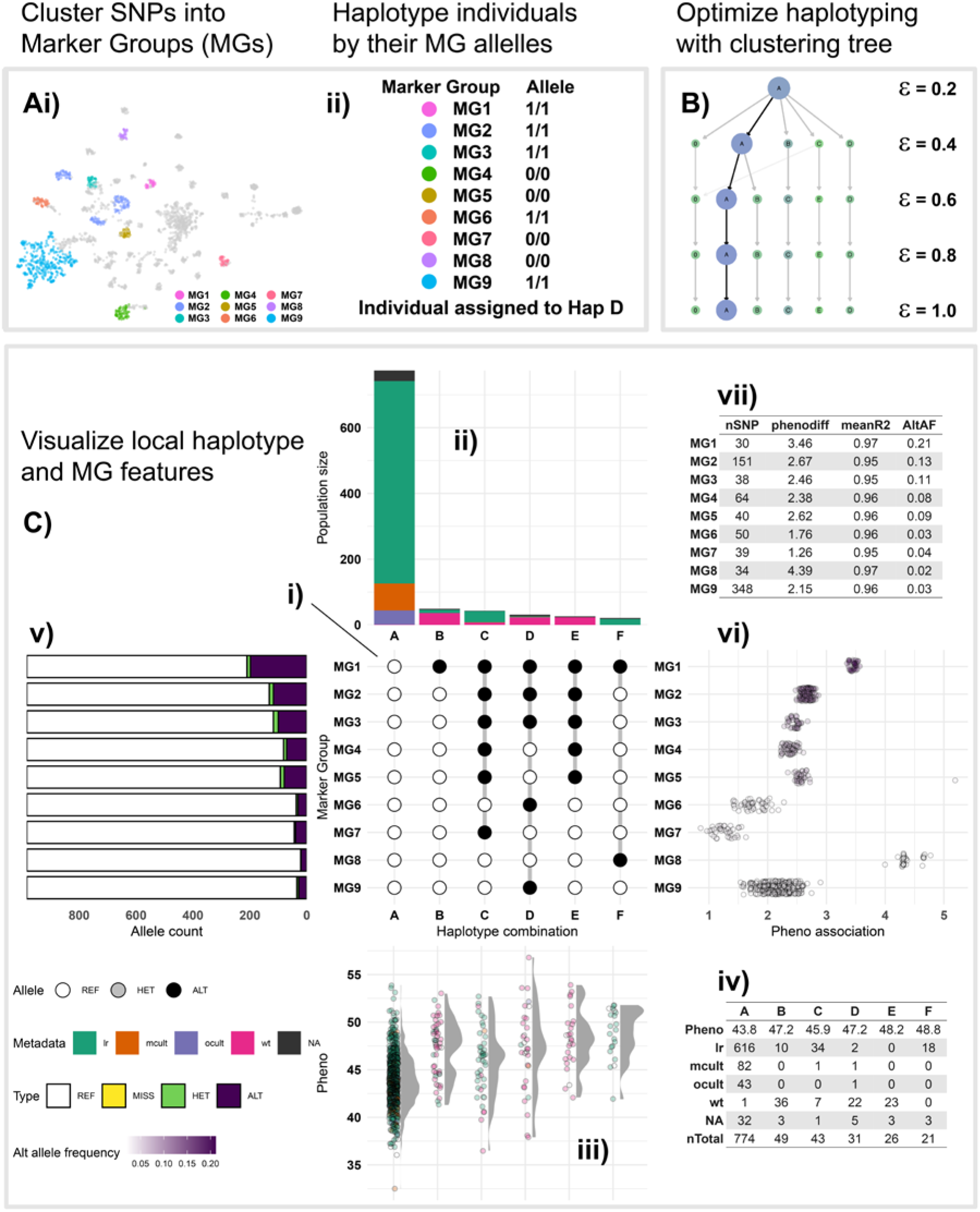
Overview of the local haplotype analysis pipeline performed by the three core ‘crosshap’ functions. ‘run_haplotyping’ performs Marker Group generation, represented by a UMAP of all SNPs co-located based on linkage, coloured by assigned Marker Group, (**Ai**), as well as haplotyping of individuals (**Aii**), before returning haplotype objects with results. ‘clustree_viz’ generates a clustering tree (**B**) to aid users in choosing an optimal epsilon value. ‘crosshap_viz’ generates a comprehensive visualization of a haplotyping results (**C**). The Marker Group alleles defining each haplotype are represented in the centre (**Ci**), results related to individuals and haplotypes are visualized vertically (**Cii**-**iv**), results related to Marker Groups and SNPs are visualized horizontally (**Cv**-**vii**).

### 2.2 Defining haplotypes

The first stage of local haplotyping with ‘crosshap’ is clustering of SNPs into representative ‘Marker Groups’ (MGs) using density-based spatial clustering of applications with noise (DBSCAN) (Ester et al., 1996) (see Supplementary File 1 for discussion of alternative variant clustering algorithms). Following this, the most common (arithmetic mode) allelic states (homozygous reference/homozygous alternate/heterozygous) across the SNPs within each Marker Group are calculated for each individual, which in turn becomes their assigned allele for the entire Marker Group (Figure 1Ai).

Following the assignment of Marker Group allelic states for each individual, the frequency of each unique allelic combination is reported. Haplotype combinations with a minimum frequency of individuals (default minHap = 9) are assigned a unique label (A-Z) and kept for downstream analysis. Marker groups that are invariant across the haplotype combinations are removed, and the remaining marker groups are provided with the prefix ‘MG’ and an identifying number (e.g. ‘MG1’). Individuals with a Marker Group allele combination corresponding to a haplotype are assigned to a haplotype group (Figure 1Aii). ‘run_haplotyping’ returns a haplotype object containing results for several user-provided epsilon values.

### 2.3 Optimization with clustering tree

To aid users in choosing an optimal epsilon value for DBSCAN clustering of variants, ‘crosshap’ provides a clustree (Zappia and Oshlack, 2018) wrapper that summarizes differences between marker groups and haplotype populations identified at different epsilon levels in a clustering tree (Figure 1B). The clustering tree (Figure 1B) fulfils the same role as the commonly used k-nearest neighbour (KNN) distance plot (Kriegel et al., 2011) by displaying cluster stability at different epsilon values. However, it is better suited to optimizing variant dimension reduction for local haplotyping because a clustering tree isolates the effects of epsilon changes on each individual cluster rather than reflecting the mean KNN distance across all points, which is sensitive to outliers. In addition to displaying cluster stability and size (nSNP or nIndividuals), the ‘crosshap’ clustering tree indicates the phenotypic association of each Marker Group or haplotype population.

## 3 Visualizing haplotypes

The ‘crosshap’ visualization provides users with a comprehensive dashboard incorporating several plots to intuitively connect marker group features with haplotype combination attributes. This allows users to quickly identify which individuals or demographic group frequently possess specific Marker Group alternate alleles, providing insight into the patterns of genomic variability across populations.

Local haplotype visualization with ‘crosshap’ is anchored around a central matrix that depicts the Marker Group allelic combinations defining each haplotype group (Figure 1Ci), modelled from the concept of UpSet plots (Lex et al., 2014). This can be useful to identify nested linkage patterns, where specific Marker Groups may be shared between haplotypes that differ across other variants. Surrounding the central matrix are peripheral plots, with attributes of the individuals in each haplotype displayed vertically in the top and bottom plots (Figure 1Cii-iii). Across each haplotype, their frequency, phenotype scores and spread of individuals from different demographic groups are captured (Figure 1Cii-iii), with additional functionality to isolate and report the phenotype association of specific demographic groups with the ‘isolate_groups’ option.

Attributes of SNPs in each Marker Group are displayed horizontally in the left and right plots (Figure 1Cv-vi). The frequencies of the different allelic states are captured as well as the phenotype association. The phenotype association of SNPs is calculated as the difference between individuals with reference and alternate alleles, assuming additive effects for heterozygous individuals (i.e. phenotype scores from homozygous individuals are weighted double those from heterozygous individuals). Alternative plots are provided to users with the ‘plot_left’ and ‘plot_right’ switches, which can be used to instead display the relative genomic positions of Marker Group SNPs across the region of interest, and a boxplot that represents the extent and sphericity of intra-Marker Group linkage. Additional tabular summaries are provided to supplement the plots with haplotype information in the bottom-left (Figure 1Civ) and Marker Group information in the top-right (Figure 1Cvii).

Detailed vignettes are provided alongside documentation to enable users to prepare inputs, learn the core functionality of ‘crosshap’, and build a UMAP GIF visualization to aid in conceptualizing the variation captured by local haplotyping at https://jacobimarsh.github.io/crosshap/).

## 4 Conclusion and Perspectives

The primary use-case of ‘crosshap’ is as a follow-up to GWAS or QTL mapping, to elucidate patterns of inheritance in a trait-associated interval of interest. Visualization with ‘crosshap’ aids in characterizing genomic variation in the region, and is valuable for uncovering novel features and relationships between haplotypes. Following this, clusters of SNPs defined in Marker Groups and the haplotype assignments of individuals become useful for enabling downstream population genetics analysis between relevant groups.

The development of ‘crosshap’ represents a major advance in the accessibility of reproducible local haplotyping tools. With a focus on guiding parameter optimization and intuitive visualizations that connect genomic variation to individual-level traits, ‘crosshap’ makes robust local haplotyping accessible to users with limited domain knowledge, equipping users with the tools necessary to develop a deep understanding of linkage patterns in a genomic region of interest.

As trait mapping and association analysis moves toward the widespread use of unsupervised learning approaches with ‘black box’ issues of interpretability, it will be increasingly necessary to follow up on these results by effectively visualizing patterns of local genomic variation to translate the results to the user. Capturing complex linkage structures and local haplotype combinations is an essential component for understanding the inheritance of variants in a genomic region of interest, a task that is made more accessible with the development of ‘crosshap’.

## Supplementary data

Supplementary data are available at *Bioinformatics* online.

## Data availability

Data used to generate the figure is available at figshare (https://figshare.com/articles/dataset/fin_b51_173kb_only_vcf/22331167) and is made available as example data within the ‘crosshap’ package.

## Supplemental 1: Variant dimension reduction clustering algorithms

The aim of variant dimension reduction for local haplotyping is to identify distinct groups of co-inherited variants in high LD with each other that can be represented by single data points, which are termed ‘marker groups’. The fewer the number of groups that variants are clustered into, the fewer the dimensions of variability which in turn improves interpretability of the haplotypes.

However, decreasing dimensionality too far comes at the cost of internal marker group homogeneity and leads to highly dissimilar variants being treated as identical in downstream analysis. To maximize the utility of each marker group, crosshap’s variant clustering only retains core groups of variants that are highly internally linked, whilst removing independent variants that are not part of major linkage patterns in the region (see main text).

### Centroid-based clustering

Local haplotyping involves characterizing high-dimensional variant data for which the correct number of groups to assign variants cannot be known or estimated without complete ARG information (local trees needed), especially as overall genealogical relatedness between individuals and populations will not necessarily be directly reflected in local genomic heritability patterns. As a result, traditional centroid-based clustering methods such as K-means and K-mediods that rely on users to define the number of clusters in a dataset *a priori* are not ideally suited for the task of variant clustering (Thalamuthu et al., 2006; Karim et al., 2021). X-means is a centroid-based alternative that overcomes this limitation by automatically determining the number of clusters. However, as with other centroid-based methods, X-means relies on the assumption that there is a central node for each cluster that correlated variables are placed around in a spherical Gaussian distribution (Tomasev et al., 2014). This is inappropriate for clustering variants as complex population structure, selection effects on distal unlinked QTLs and recombination stochasticity are likely to result in linked marker groups that are not spherically distributed around a central point, even if a single variant has experienced direct selection.

### Hierarchical clustering

Hierarchical clustering improves over basic centroid-based methods by aiding the user in selecting the optimal number of clusters (Rokach and Maimon, 2005). However, both hierarchical and centroid-based methods are highly sensitive to outliers, and are not intended to identify outliers, as they primarily work by partitioning all data points based on their relationships with all other data points (Karim et al., 2021). For the purposes of variant dimension reduction, this dilutes the effect of local clusters of densely linked variants, which are the patterns that need to be identified and isolated.

### Density-based clustering

Density-based spatial clustering of applications with noise (DBSCAN) (Ester et al., 1996) is an algorithm implemented for variant dimension reduction in crosshap. DBSCAN is well-suited to high-dimensional variant data and possesses several key advantages over hierarchical and centroid-based clustering for capturing linked marker groups (Karim et al., 2021). Firstly, DBSCAN defines clusters by only considering the density of proximal (correlated) points and classifies points that are not in a dense cluster of points as noise. Therefore, not only does DBSCAN natively bypass the need for applying post-hoc outlier detection algorithms, but more critically, the clusters themselves are defined without being biased by noise in the dataset (Schubert et al., 2017).

DBSCAN ‘builds’ clusters from nodes of dense points by finding connected paths with other densely correlated neighbouring points (Ester et al., 1996). Therefore, DBSCAN does not assume cluster sphericity and can classify marker groups that are irregularly distributed. However, a drawback of this feature is that when large clusters are non-spherical, distant points within the cluster can be unpredictably dissimilar to each other; i.e., there is no theoretical limit to how uncorrelated two points within a cluster can be if there are enough intermediate points ‘connecting’ them. This pitfall is overcome in crosshap by a ‘smoothing’ step during variant dimension reduction that removes outlier points in each cluster.

### Variant cluster smoothing

The cluster ‘smoothing’ is performed by calculating the mean pairwise linkage (R^2^) of each SNP in a cluster with all other points within the same cluster. Loci that are in very high linkage with many other points in the cluster will have a high mean intra-cluster R^2^, whereas outlier loci will have a low mean intra-cluster R^2^. To classify outliers, the standard deviation of the mean intra-cluster R^2^ is calculated, and loci that exhibit a score 2 standard deviations below the median of the mean intra-cluster R^2^ for a given cluster will be removed (Supplemental Figure 1). Cluster ‘smoothing’ ensures that while initial clustering is performed without assumptions regarding the shape of the data, clusters are marginally pruned to ensure that each locus within a marker group is in high linkage with all other loci, removing the number of outlier SNPs (Supplemental Figure 1). The ‘smoothing’ establishes a minimum level of sphericity between SNPs within each marker group to improve internal homogeneity (Supplemental Figure 1B).

**Supplemental Figure 1.**
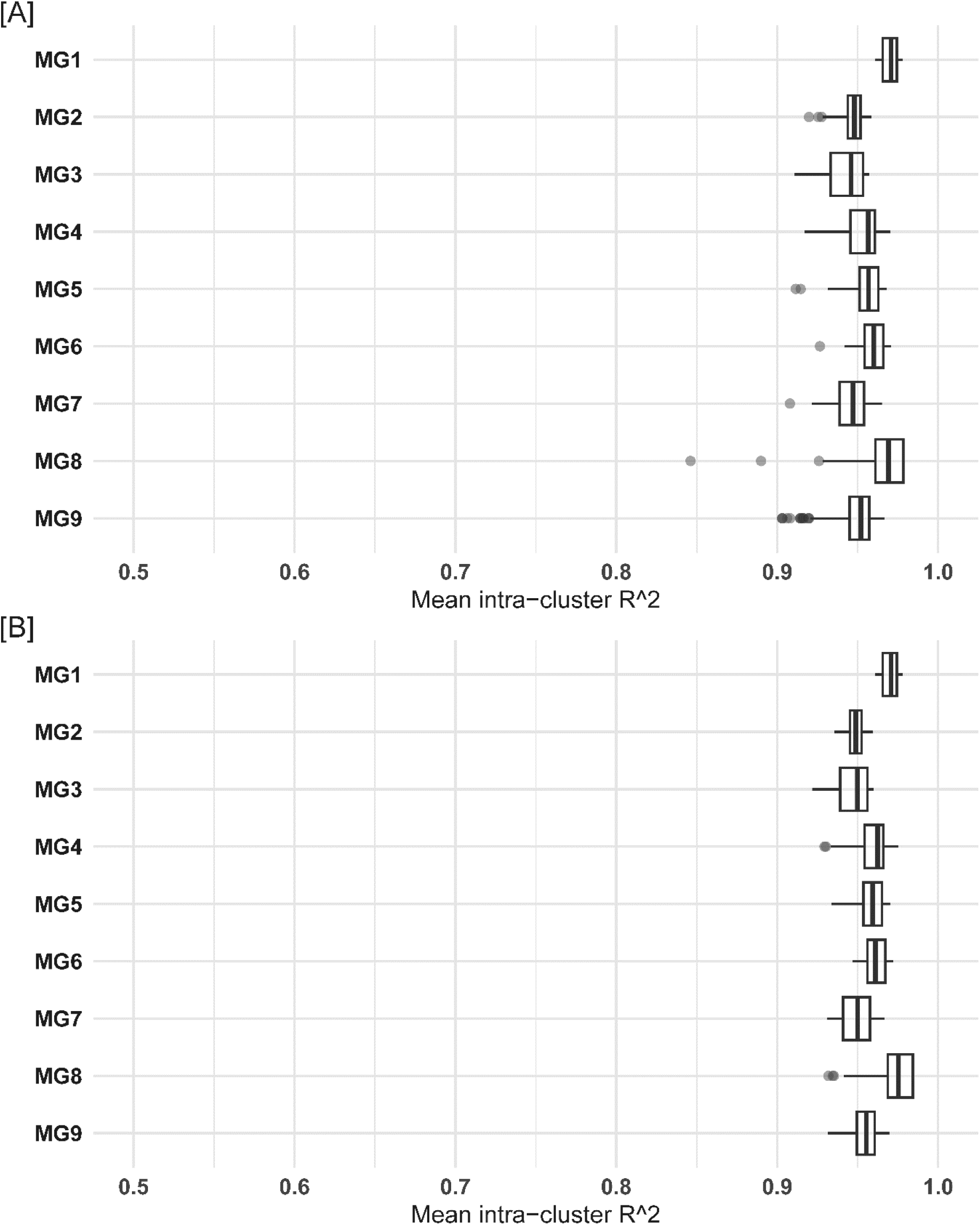
Boxplot of mean linkage (R^2^) of each SNP with other SNPs within the same marker group; without cluster smoothing **[A]** and after cluster smoothing **[B]**. N.B. outlier points in **[B]** are defined in reference to the mean and standard deviation of the marker groups after smoothing and were not greater than 2 standard deviations below the median in **[A]**.

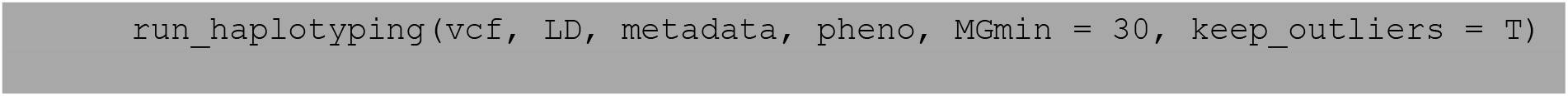
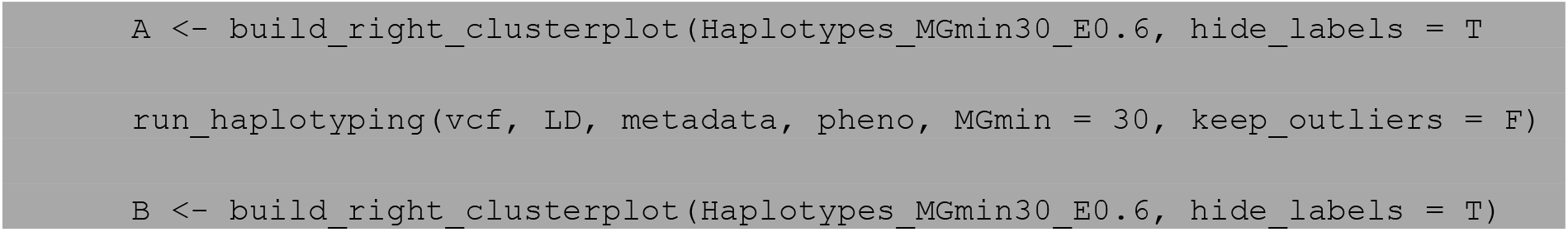

Successful clustering with DBSCAN is highly contingent on the use of appropriate minimum points (minPts) and epsilon parameters. In the context of local haplotyping, minPts may be set based on the minimum number of SNPs the user chooses to allow in each marker group. This may change depending on the density of SNPs captured in the region, though values in the range of 10-30 will typically be appropriate. Epsilon is a statistical term that in the case of DBSCAN, refers to the local radius of points in a cluster within which new points are counted and recursively incorporated (if they exceed the minPts threshold). Choosing an appropriate epsilon value is a major obstacle to the accessibility of local haplotyping using DBSCAN, as even though epsilon roughly translates to ‘cluster density’, it is highly dataset dependent and laborious to optimize (Kriegel et al., 2011).

### Hierarchical density-based clustering

Hierarchical density-based spatial clustering of applications with noise (HDBSCAN) (McInnes et al., 2017) is an extension of DBSCAN which bypasses the need to specify an epsilon parameter. While DBSCAN requires a static epsilon value across an entire dataset, HDBSCAN instead algorithmically optimizes the epsilon parameter separately for each cluster. Therefore, HDBSCAN can capture clusters of varying densities, and only requires a minPts parameter as input. A (now deprecated) HDBSCAN implementation of local haplotyping is available in crosshap and was tested against a carefully chosen DBSCAN epsilon parameter (0.6).

**Supplemental Figure 2.**
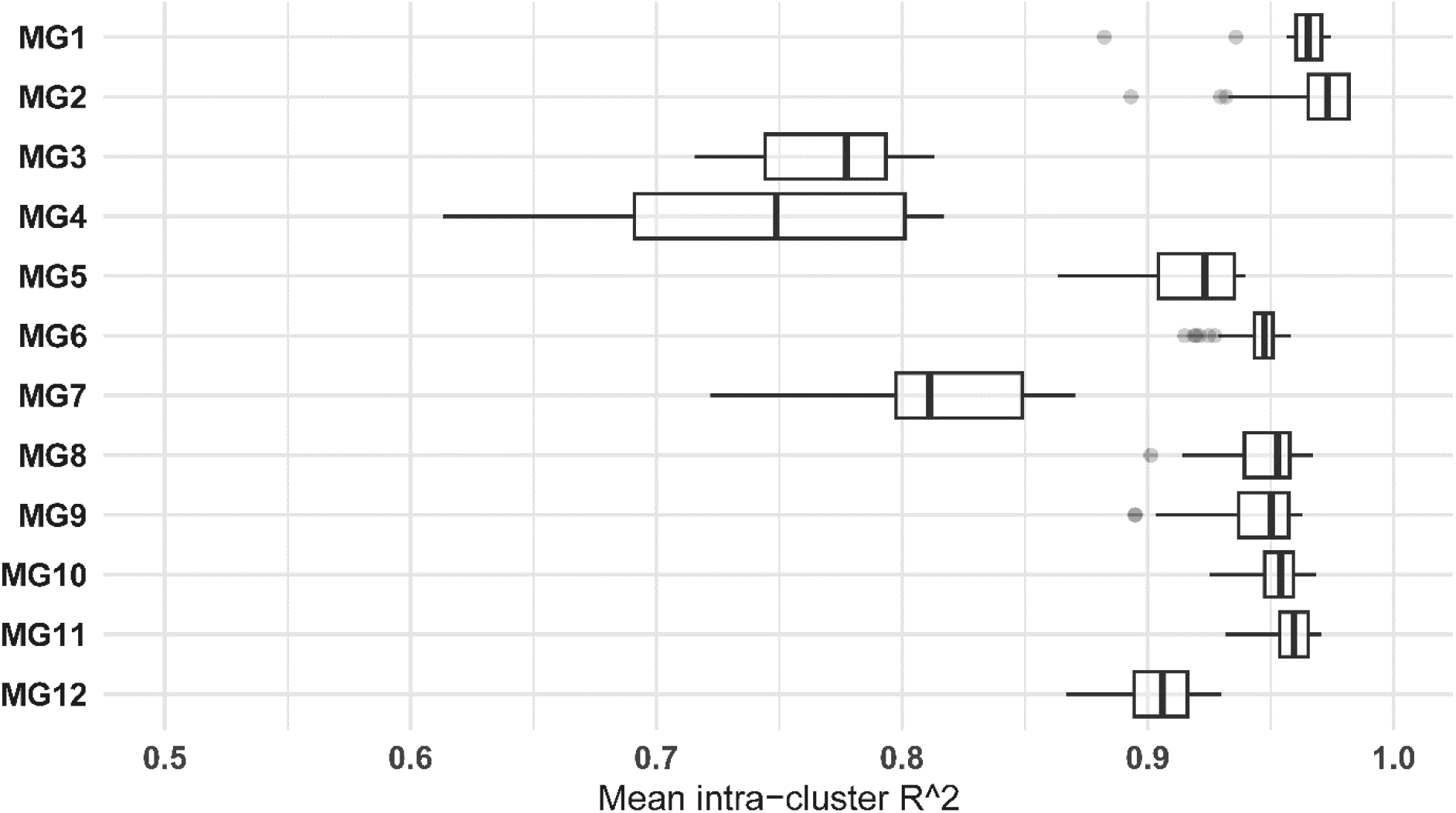
Boxplot of mean linkage (R^2^) of each SNP with other SNPs within the same marker group defined by HDBSCAN with cluster smoothing.

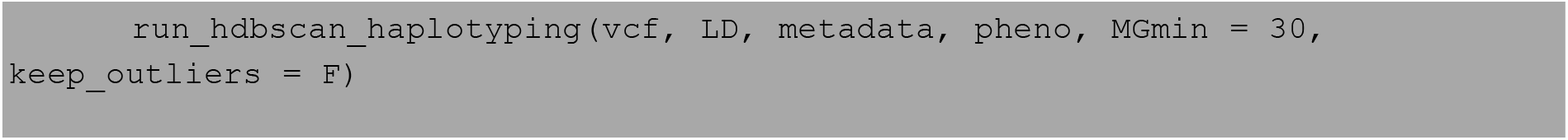
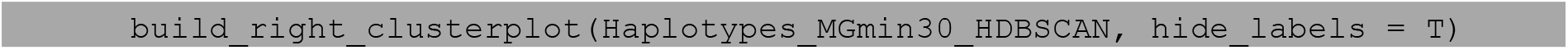

HDBSCAN identified many marker groups; however, this led to the clustering of marker groups that possessed poor internal linkage (Supplemental Figure 2). Variability in the density of clusters is not necessary for local haplotyping as all marker groups need to be highly internally linked, which can be provided by a single stringent epsilon value. Therefore, DBSCAN was chosen as the default clustering algorithm for variant reduction and tools were developed to aid the user in choosing an epsilon parameter that is optimal for a given dataset (see main text).

## References

Belzile F, Abed A, Torkamaneh D (2020) Time for a paradigm shift in the use of plant genetic resources. Genome 63: 189–194

Ester M, Kriegel H-P, Sander J, Xu X (1996) A density-based algorithm for discovering clusters in large spatial databases with noise. In Proceedings of the 2nd ACM International Conference on Knowledge Discovery and Data Mining (KDD), pp 226–231

Kriegel HP, Kröger P, Sander J, Zimek A (2011) Density-based clustering. WIREs Data Mining and Knowledge Discovery 1: 231–240

Lex A, Gehlenborg N, Strobelt H, Vuillemot R, Pfister H (2014) UpSet: Visualization of Intersecting Sets. IEEE Transactions on Visualization and Computer Graphics 20: 1983–1992

Li X, Shi Z, Gao J, Wang X, Guo K (2023) CandiHap: a haplotype analysis toolkit for natural variation study. Molecular Breeding 43

Marsh JI, Hu H, Petereit J, Bayer PE, Valliyodan B, Batley J, Nguyen HT, Edwards D (2022) Haplotype mapping uncovers unexplored variation in wild and domesticated soybean at the major protein locus cqProt-003. Theor Appl Genet 135: 1443–1455

Marsh JI, Petereit J, Monahan G, Edwards D, Bayer PE (2022) Recombination, linkage and haplotypes: dissecting patterns for inheritance for genetic gain, Vol 12. CABI Biotechnology, Oxfordshire, UK

Pruim RJ, Welch RP, Sanna S, Teslovich TM, Chines PS, Gliedt TP, Boehnke M, Abecasis GR, Willer CJ (2010) LocusZoom: regional visualization of genome-wide association scan results. Bioinformatics 26: 2336–2337

Schaid DJ, Chen W, Larson NB (2018) From genome-wide associations to candidate causal variants by statistical fine-mapping. Nat Rev Genet 19: 491–504

Tardivel A, Torkamaneh D, Lemay MA, Belzile F, O’Donoughue LS (2019) A systematic gene-centric approach to define haplotypes and identify alleles on the basis of dense single nucleotide polymorphism datasets. Plant Genome 12

Wang CC, Yu H, Huang J, Wang WS, Faruquee M, Zhang F, Zhao XQ, Fu BY, Chen K, Zhang HL, Tai SS, Wei C, McNally KL, Alexandrov N, Gao XY, Li J, Li ZK, Xu JL, Zheng TQ (2019) Towards a deeper haplotype mining of complex traits in rice with RFGB v2.0. Plant Biotechnology Journal 18: 14–16

Wu X, Jiang W, Fragoso C, Huang J, Zhou G, Zhao H, Dellaporta S (2022) Prioritized candidate causal haplotype blocks in plant genome-wide association studies. PLoS Genet 18

Zappia L, Oshlack A (2018) Clustering trees: a visualization for evaluating clusterings at multiple resolutions. Gigascience 7

## References

Karim MR, Beyan O, Zappa A, Costa IG, Rebholz-Schuhmann D, Cochez M, Decker S (2021) Deep learning-based clustering approaches for bioinformatics. Brief Bioinform 22: 393–415

Kriegel HP, Kröger P, Sander J, Zimek A (2011) Density-based clustering. Wiley Interdiscip Rev: Data Min Knowl Discov 1: 231–240

McInnes L, Healy J, Astels S (2017) hdbscan: Hierarchical density based clustering. J Open Source Softw 2

Rokach L, Maimon O (2005) Clustering Methods. In Data Mining and Knowledge Discovery Handbook, pp 321–352

Schubert E, Sander J, Ester M, Kriegel HP, Xu X (2017) DBSCAN Revisited, Revisited. ACM Transactions on Database Systems 42: 1–21

Thalamuthu A, Mukhopadhyay I, Zheng X, Tseng GC (2006) Evaluation and comparison of gene clustering methods in microarray analysis. Bioinformatics 22: 2405–2412

Tomasev N, Radovanovic M, Mladenic D, Ivanovic M (2014) The Role of Hubness in Clustering High-Dimensional Data. IEEE Transactions on Knowledge and Data Engineering 26: 739–751

